# *Zfp36l1* establishes the high affinity CD8 T cell response by directly linking TCR affinity to cytokine sensing

**DOI:** 10.1101/2023.05.11.539978

**Authors:** Georg Petkau, Twm J. Mitchell, Marian Jones Evans, Louise Matheson, Fiamma Salerno, Martin Turner

## Abstract

How individual T cells compete for and respond to IL2 at the molecular level, and, as a consequence, how this shapes population dynamics and the selection of high affinity clones is still poorly understood. Here we describe how the RNA binding protein ZFP36L1, acts as a sensor of TCR affinity to promote clonal expansion of high affinity CD8 T cells. As part of an incoherent feed forward loop ZFP36L1 has a non-redundant role in suppressing negative regulators of cytokine signalling and mediating a selection mechanism based on competition for IL2. We suggest that ZFP36L1 acts as a sensor of antigen affinity and establishes dominance of high affinity T cells by installing a hierarchical response to IL2.

## Introduction

Upon antigen encounter T cells interpret multiple cues including co-stimulatory molecules and pro- and anti-inflammatory cytokines and translate these into molecular programs which lead to clonal expansion and differentiation^1^. A diverse repertoire of T cells with varying affinities for antigen, and thus different requirements for activation, compete amongst each other for antigen, costimulation, cytokines and nutrients^2,3^. The outcome of this competition is that few high affinity T cell clones dominate at the peak of the T cell response. Multiple mechanisms have been proposed to control the expansion and differentiation of an antigen specific CD8 T cell population^4,5^. These include the frequency of homotypic cell contacts determined by cell density and the consequent impact of inhibitory receptors expressed by the expanding population^4,5^. In addition, intercellular communication via local gradients of cytokines such as IL2 has been suggested to be essential to adjust the clonal sizes of effector and memory populations^4,6^. Access to IL2 can be licensed by other inflammatory cytokines including type-I interferons and IL12 which enhance clonal expansion and effector differentiation^7,8^. It has also been shown that strongly activated T cell clones are able to enhance the proliferation of weakly activated clones by the production of an excess of IL2, at least *in vitro*^9^. Using expression of Nur77-GFP as a reporter for TCR signaling, a threshold for activation has been suggested to induce cell proliferation and to be invariable to different, above-threshold, stimulation strengths^10^. Costimulatory cues, including IL2, can lower cumulative activation thresholds of T cells whereby these signals are integrated by transcription factors which drive metabolic reprogramming, proliferation and differentiation ^11–13^. Interestingly, the timing of initiation of cell division, acquisition of effector function and the underlying transcriptional pathways were not dependent on CD8 T cell affinity^14^. In line with this, stimulation of naïve OT-I cells with lower affinity antigens *in vivo*, did not result in differential initial expansion of T cells early in the response^15^. High affinity T cells establish their larger clone size rather by additional rounds of cell division^15^. In the later phase of the immune response T cell proliferation is driven by IL2 and the prolonged expression of CD25, promoted by inflammatory cytokines^8,16^. In addition, inflammation has been suggested to rescue high affinity but not low affinity T cells from apoptosis when antigen is limited^17^. Currently we are lacking a mechanistic understanding of how T cell affinity is translated into differential response to cytokines and preferential expansion of high affinity clones.

Autocrine and paracrine IL2 are to some extent redundant as autocrine IL2 is largely dispensable for primary CD8 T cell expansion *in vivo*^18–20^, but its production by other cells is essential ^16,19,21–23^. Access to IL2 is critical to regulate apoptosis in a TCR dependent fashion resulting in the preferential survival of high affinity clones^24^. Thus competition for IL2 by different T cell clones and subsets may be essential to shape the T cell immune response and the establishment of dominance by high affinity T cell clones^2,25–27^. It is to date not clear how T cells of different affinities which are recruited into the immune response and thus have overcome thresholds of activation establish the duration of the response to IL2. How TCR affinity is directly linked to the sensitivity to IL2 during an immune response remains unknown.

The ZFP36 family of RNA binding proteins are best characterised for their role as suppressors of cytokine production. Three paralogues (*Zfp36, Zfp36l1* and *Zfp36l2*) are expressed by T cells where they have been shown to limit cytokine production, and more recently, to regulate metabolic and transcriptional programs ^28–31^. CD8 T cells lacking both *Zfp36* and *Zfp36l1* displayed enhanced *in vitro* expansion, accelerated differentiation, and more potent effector function. Moreover, ZFP36 and ZFP36L1 enforced dependence on CD28 costimulation. ZFP36L2 has a unique role in repressing cytokine production by memory T cells^28^, but because of an apparent redundancy between the *Zfp36* paralogues^29–31^ unique roles for individual ZFP36 family members in T cell activation and differentiation have yet to be identified. In the present study we show that ZFP36L1 acts as a node which senses TCR-ligand affinity and promotes sensitivity to cytokine signals, thereby organizing inter-clonal competition and safeguarding the selection of high affinity T cell clones.

## Results

### ZFP36 and ZFP36L1 confer a competitive advantage to CD8 T cells in vitro

Naïve CD8 T cells from OT-I TCR transgenic (TCR specific for the SIINFEKL peptide from chicken ovalbumin) CD4^cre^ ZFP36^fl/fl^ ZFP36L1^fl/fl^mice, lacking *Zfp36* and *Zfp36l1* in T cells (from now on termed dKO) proliferate more extensively than OT-I cell isolated from CD4^wt^ ZFP36^fl/fl^ZFP36L1^fl/fl^littermate controls, after *in vitro* stimulation with high affinity SIINFEKL (N4) peptide. When cultured alone they accumulate substantially greater numbers of cells that had undergone more cell divisions after 48h (**Fig.1a**). Since both RBPs have been previously shown to limit autocrine cytokine production including IL2^30,31^, which could feedback as costimulatory signals we wanted to evaluate whether the greater expansion is due to increased production of cytokines by the dKO. Therefore, we performed co-culture experiments where both cell types are exposed the same environment and can interact with each other. In co-cultures after 48h of stimulation, expanding WT cells showed a similar distribution of cells per generation as the dKO but accumulated more cells per generation (**Fig.1b**). Thus, when exposed to the same environment, dKO cells lost their superior ability to expand.

WT cells co-cultured with dKO cells accumulated greater cell numbers than the dKO cells and these numbers exceeded those of WT cells cultured alone (**Fig.1c**). Similar results were obtained with non-transgenic naïve CD8 T cells stimulated with anti-CD3 and anti-CD28 (**Supplementary Figure 1a, b)**. The proliferation index, a measure of cell intrinsic capacity to divide, of WT cells is increased when co-cultured with dKO cells and reaches similar levels as single cultured dKO cells **(Fig.1d)**. The cell division of the dKO cells is only very slightly reduced upon co-culture, suggesting the reduced numbers of dKO cells in the presence of WT cells reflects increased cell death. Thus, when WT and dKO cells were cultured in the same well, the dKO cells were less able to compete with their WT counterparts.

**Figure 1:**
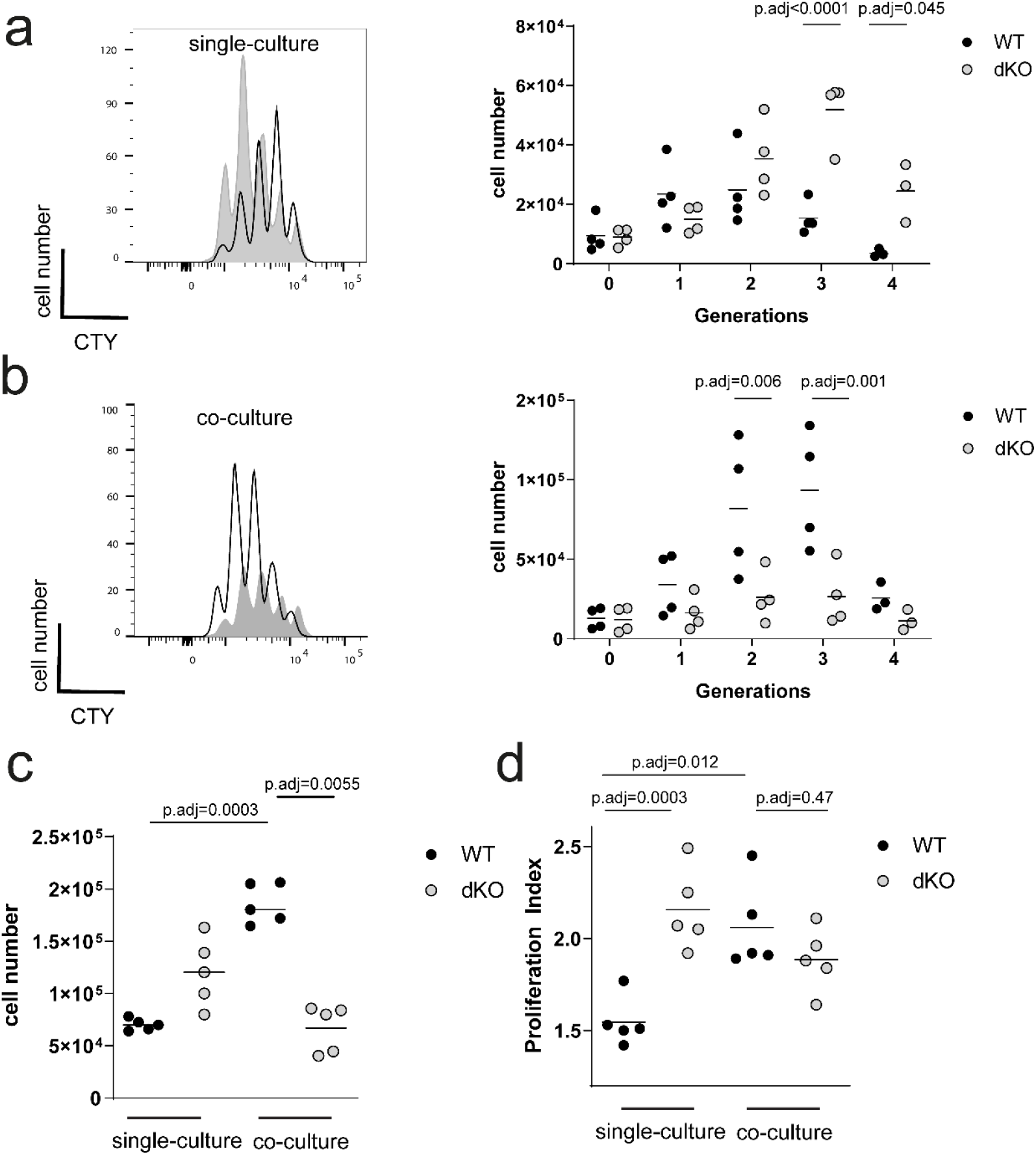
ZFP36 and ZFP36L1 confer a competitive advantage to CD8 T cells in vitro. **a, b)** Left panels show the dilution of CTY by naïve WT (open) and dKO (filled) OT-I cells stimulated with N4 peptide for 48 hours individually **a**) or in co-culture **b**). Right panels show the absolute cell numbers per generation in single **a**) and co-cultures **b)**: Data is compiled from 4 independent experiments. Each data point is representative of one biological replicate. **c)** Absolute cell numbers of WT and dKO OT-I cells after 48 hours with N4 peptide in individual and co-culture. Data is compiled from 5 independent experiments. **d)** Proliferation index of cells as in panel **c)**. Statistical significance in **a, b**) was determined by 2-way ANOVA followed by Bonferroni multiple comparisons test. Statistical significance in **c, d)** was determined by one-way ANOVA followed by Tukey’s multiple comparisons test. Cell numbers in co-cultures are adjusted to the double culture volume to compare to WT culture conditions, maintaining the same cell culture density. The statistical mean is indicated by horizontal line in **a, b)**.

### RBP promote T cell fitness by enhancing the response to IL2

To assess the response to IL2, which is critical for T cell survival and proliferation *in vitro*, we activated WT and dKO OT-I cells with high affinity N4 peptide for 48 hours alone or in co-culture. When cultured alone WT cells expressed low levels of CD25, the high affinity subunit of the IL2 receptor, when compared to dKO cells. Upon co-culture CD25 was increased on WT cells, but its expression was reduced on dKO cells compared to single cultures (**Fig. 2a, b**). The greater expression of CD25, when dKO cells are cultured alone, likely reflects the increased amount of IL2 produced since the expression of CD25 can be induced by IL2^32^. When in co-culture and exposed to similar amounts of IL2, CD25 expression increases (**Fig.2b**) and remains higher on WT cells compared to dKO, which suggests decreased IL2 signalling in dKO cells.

**Figure 2:**
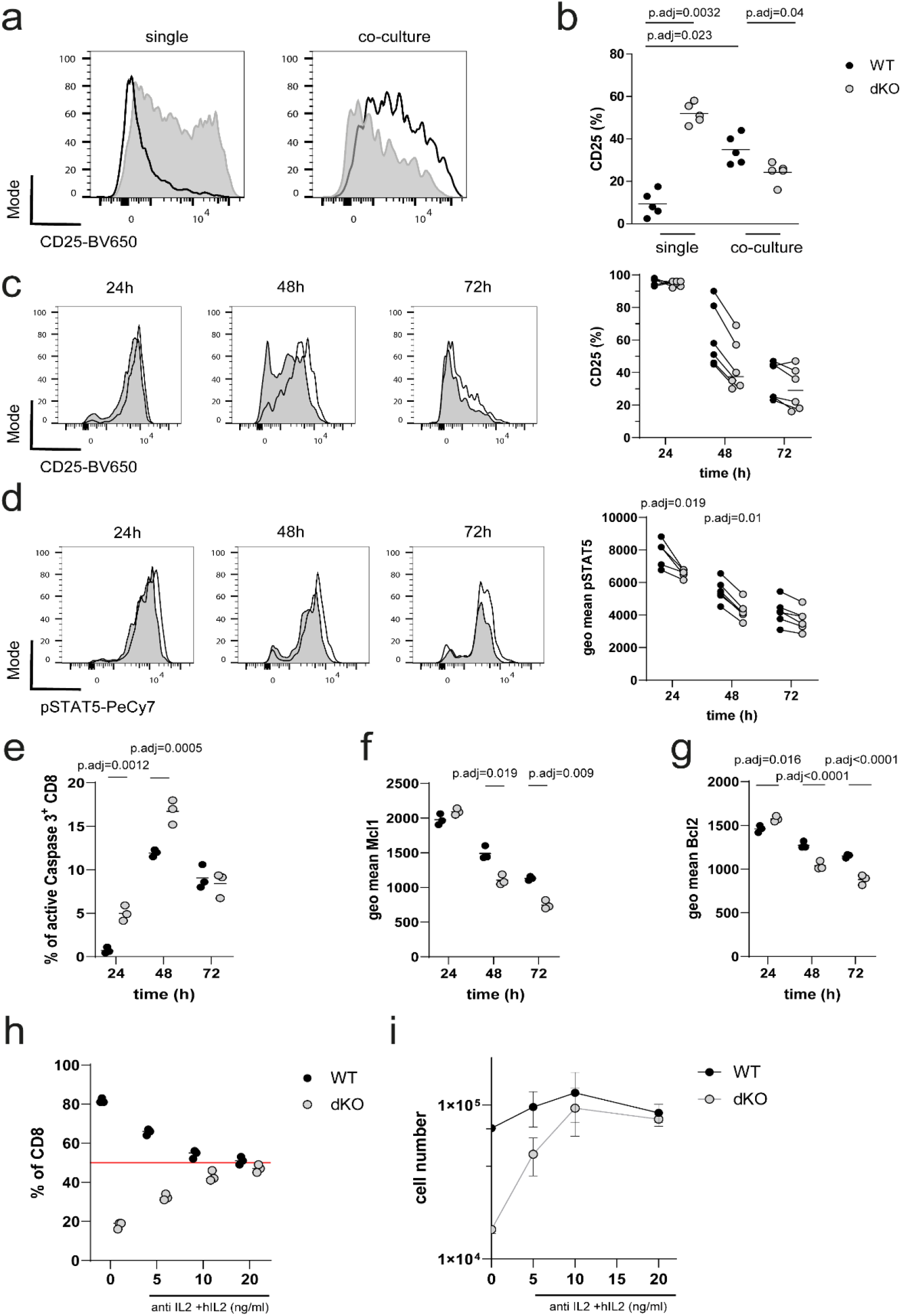
RBP promote T cell fitness by enhancing the response to IL2. **a)** Representative histograms showing CD25 surface expression in WT (open) and dKO (filled) OT-I cells in single and co-cultures, 48h hours after N4 peptide stimulation. **b)** Frequency of CD25^+^ WT and dKO OT-I cells in single and co-cultures. Each data point represents one biological replicate. Data is compiled from five independent experiments. **c)** Left panel shows time course analysis of intracellular CD25 expression as representative histograms of cocultured WT and dKO OT-I cells, which were stimulated with N4 peptide for indicated times. Right panel shows frequency of CD25 positive cells. **d)** Left panel shows representative histograms of pSTAT5 over indicated times after N4 peptide stimulation of WT and dKO co-cultured OT-I cells. Right panel show pSTAT5 the geo mean expression per cell. Statistical significance was determined by two-way ANOVA followed by Bonferroni test for multiple comparisons. Data is compiled from 2 out of three independent experiments with each data point representing a technical replicate. **e)** Frequencies of active-CASPASE3 positive OT-I cells in co-cultures stimulated with N4 peptide. Geo mean fluorescence intensity of **f)** Mcl1 and **g)** Bcl2 in co-cultured WT and dKO OT-I cells is shown. Data is representative of two in e) and three in f, g) independent experiments with each data point representing a technical replicate. Statistical significance was determined by one-way ANOVA followed by Tukey’s test for multiple comparisons. Frequencies **h)** and absolute cell numbers **i)** of WT and dKO co-cultured OT-I cells, 72h after stimulation with N4 peptide in normal medium or in the presence of anti-mouse IL2 antibody and indicated amounts of added human IL2 from the beginning of the culture. Data is representative of two independent experiments with each data point representing a technical replicate. The statistical mean is indicated by horizontal line in all panels. Error bars indicate the SD of the mean in all panels.

Next, we assessed the dynamics of IL2 responsiveness following antigen activation, by staining of intracellular CD25 and phosphorylated pSTAT5 (pY694) as a marker of IL2 signal transduction in co-culture. 24h after activation the frequencies of CD25^+^ cells were the same between WT and dKO, and pSTAT5 was only slightly reduced in dKO (**Fig.2c, d**). At 48h after activation, both the frequency of CD25^+^ cells and the phosphorylation of STAT5 were reduced in the dKO compared to WT OT-I (**Fig.2c, d**). There was a higher frequency of active CASPASE3-positive dKO cells than WT cells 24 and 48h after activation (**Fig.2e**). This was accompanied by a reduced expression of the anti-apoptotic proteins MCL1 and BCL2 in dKO cells compared to the WT cells, at later time points following activation (**Fig.2f, g, Supplementary Figure 1c, d**). The increased apoptosis in dividing dKO cells may reflect diminished STAT5 activity and lesser amounts of antiapoptotic proteins.

The reduced expression of CD25 and decreased signalling via STAT5 in dKO cells in coculture, prompted us to suspect that dKO cells are inferior in processing IL2 signals when in competition with WT cells. We therefore controlled the concentration of IL2 present in the cultures using anti-mouse IL2 blocking antibody (JES6-1A12), while providing different concentrations of human IL2 (hIL2) which is not bound by JES6-1A12. Adding increasing amounts of hIL2 normalized the ratios of WT and dKO, reaching 50% when hIL2 was added at 20ng/ml (**Fig.2h**). The absolute cell numbers recovered from these cultures 72h after activation were also comparable between WT and dKO at high doses of hIL2 (**Fig.2i**). This data shows that when amounts of IL2 are low the survival of the dKO cells is compromised due to reduced IL2 responsiveness.

### Competitive fitness for cytokines is controlled by RBPs during bacterial infection *in vivo*

To examine the fitness of dKO cells in a competitive setting *in vivo*, we transferred 500 WT (CD45.1^+^) and dKO (CD45.2^+^) naive OT-I cells into recipients in which the host cells were positive for both CD45.1 and CD45.2. In parallel, to generate controls with the same number of high affinity antigen specific cells, we transferred 1000 WT or dKO naïve OT-I cells into separate hosts. We followed the expansion of donor OT-I cells in the blood after infection with an attenuated strain of Δ*Act Listeria monocytogenes* expressing chicken Ovalbumin (OVA_257-264_ (N4)) (*attLm*-OVA). During the expansion phase of the primary response individually transferred WT and dKO OT-I cells showed comparable frequencies in blood in contrast to the *in vitro* proliferation results. When co-transferred, the frequencies of WT OT-I cells were increased by the presence of dKO cells (**Fig.3a-c)**. Moreover, the proportion of dKO OT-I cells in blood was reduced compared to co-transferred WT OT-I cells indicating the dKO cells expand less when in competition (**Fig.3a-c**). Furthermore, the numbers of dKO OT-I cells were diminished when co-transferred with WT OT-I compared to single dKO OT-I cell transfers at the peak of the response and during contraction (**Fig.3a-c**). The diminished ability of dKO cells to compete was strikingly apparent at day 13 when dKO cells are 20-fold less frequent than their WT counterparts.

Following co-transfer of WT and dKO OT-I cells and subsequent infection with *attLm*-OVA, the frequencies of WT OT-I cells expressing CD25 on day five post infection, were higher than that of dKO cells in the spleens of recipient mice (**Fig. 3d, e**). Furthermore, the frequency of Ki-67^+^ cells, a marker of proliferating cells, was reduced amongst dKO cells compared to WT cells (**Fig. 3f, g**). Expression of CD25 and IL2 signalling extend the time of proliferation of CD8 T cells during the later phase of the primary response^8,16,22^. In line with this, our data suggests that the dKO cells exit cell cycle earlier *in vivo*. Maintenance of CD25 expression and proliferation at this stage has been attributed to the action of type I Interferons and IL12^8^. The addition of IL12 to WT and dKO OT-I cells cultured *in vitro* increased CD25 expression and only partially restored the ability of dKO cells to expand in competition (**Supplementary Figure 2a, b**). Thus, the *in vivo* and *in vitro* presence of inflammatory cues are insufficient to compensate the cell intrinsic expansion deficit of T cells lacking RBPs.

**Figure 3:**
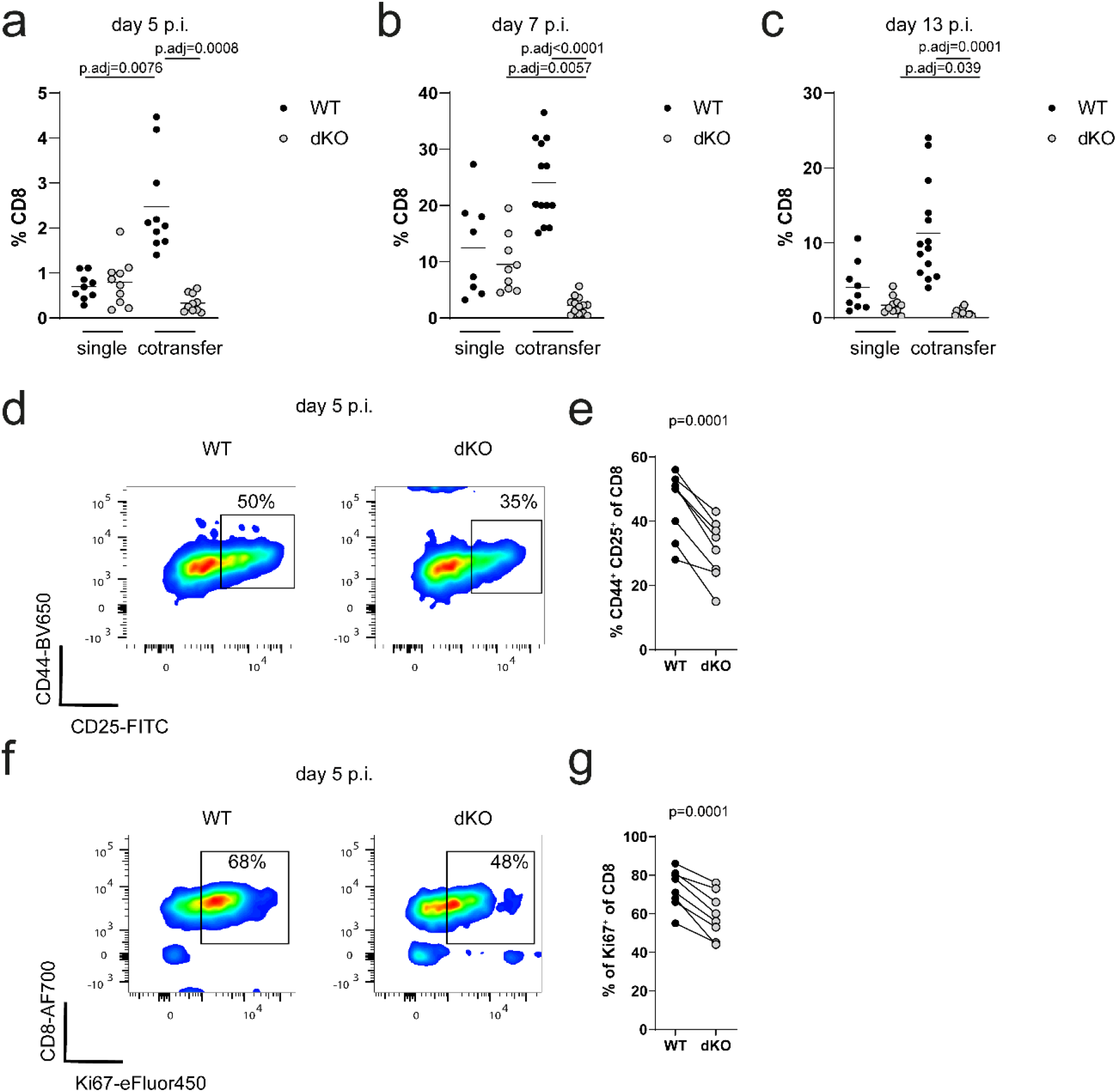
Competitive fitness for cytokines is controlled by RBPs during bacterial infection *in vivo*. **a)** Frequencies of WT and dKO OT-I cells single or co-transferred in blood of recipients on day 5,7 **b)** and 13 **c)** post infection with *attLm*-OVA. Data is pooled from three independent experiments. Statistical significance was determined by two-way ANOVA followed by Tukey’s test for multiple comparisons. **d)** Representative FACS plots showing expression of CD44 and CD25 on co-transferred WT and dKO OT-I cells in spleens of recipient mice on day 5 post *attLm*-OVA infection. **e)** Frequency of CD44^+^CD25^+^ OT-I T cells. **f)** Representative FACS plots showing expression of CD8 and Ki-67 on co-transferred WT and dKO OT-I cells. **g)** Frequency of Ki67^+^ OT-I T cells. Statistical significance in panel **e)** and **f)** was determined using a paired *t-test*. The statistical mean is indicated by horizontal line in **a-c)**.

### ZFP36 and L1 suppress a network of cytokine signaling pathway regulators

To understand the mechanism by which ZFP36 and ZFP36L1 maintain responsiveness to IL2, we performed Next Generation RNA Sequencing of naïve WT and dKO OT-I cells activated for 0, 1.5, 3, 6 and 16 hours *in vitro* with N4 peptide. Differential gene expression analysis revealed several genes which were significantly increased in dKO OT-I cells (**Fig.4a**). Amongst the differentially increased transcripts, we identified those which were directly bound by the RBPs by examining a list of direct targets identified by RNA crosslink-immunoprecipitation (CLIP) in activated CD4 (panZFP36)^33^ and CD8 (ZFP36L1)^30^ T cells (**Supplementary Table 1**).

Most direct target genes were identified among genes which were differentially increased at six hours post activation, comprising more than 40% of the increased genes at this timepoint (**Fig.4a, b**). Gene set enrichment analysis of Hallmark gene sets revealed IL2-STAT5, IL6-JAK-STAT3 and TNF-NFκB signalling pathways to be most significantly enriched among increased genes at this time (**Fig.4c**). Within the positively enriched genes among the IL2-STAT5 gene set, which were also targets of the RBP, was a subset of genes involved in the regulation of the cytokine response (**Fig. 4d**), including the family of Suppressors of Cytokine Signaling, *Socs1* and *Cish*^34^ (**Fig.4e**). *Socs3* was also identified as direct CLIP target, although the increase in its expression did not reach significance (6h timepoint, log2FC=0.39, p.adj=0.06), (**Supplementary Figure 3a**). *Socs1* is known to be induced by multiple cytokines and mediates negative feedback to limit their signaling^35^. SOCS-1^36^ and -3^37^ have been shown to be induced by IL2 and to inhibit IL2 signaling by interfering with JAK-1 and -3 signal transduction. *Socs2*, which we have found to be slightly increased 6h post activation (log2FC=0.23, p.adj=0.02) but which is not a direct RBP target, has been suggested to promote IL2 signaling via degradation of SOCS-3^38^ but also to suppress STAT5 phosphorylation in CD4 T cells^39^. The role of CISH with regards to its ability to inhibit STAT5 phosphorylation remains controversial and it may play a more prominent role in inhibiting TCR signaling^40–42^. We also identified *Pim1*, encoding a kinase which negatively regulates STAT5 signaling by stabilizing SOCS-1 and -3^43^, to be a direct RBP target (**Supplementary Figure 3b**) and to be slightly increased early in dKO OT-I cells (3h timepoint, log2FC=0.32, padj=0.06). In summary we find a network of genes to be targeted by RBPs which suppress cytokine signalling (Summarized in **Fig.4f**).

**Figure 4:**
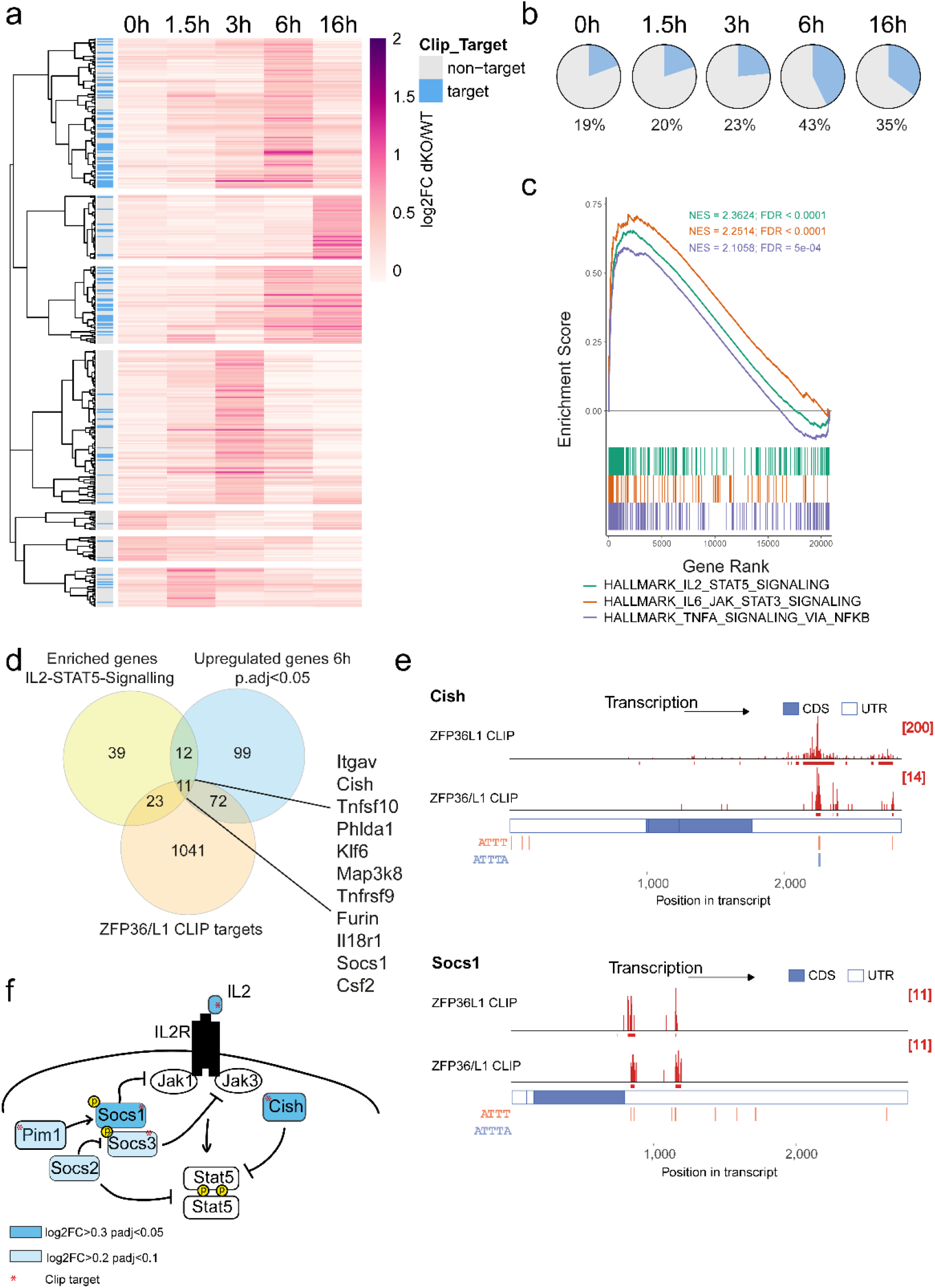
ZFP36 and ZFP36L1 suppress a network of cytokine signaling pathway regulators. **a)** Heatmap using correlation as measure of hierarchical distances, visualizes differentially upregulated genes (log2FC>0.3, adj.pvalue<0.05) between naïve WT and dKO cells at indicated timepoints after stimulation with N4 peptide. Direct CLIP target transcripts are indicated in the left column. **b)** Pie charts visualize the proportion of RBP targets among significantly differentially upregulated genes in dKO. **c)** Gene set enrichment analysis, using the murine hallmarks gene collection, shows the enrichment score versus the gene rank of the top three positively enriched gene set. FDR is indicated for each gene set. **d)** Venn diagram shows the shared genes which are contained in the positively enriched IL2-STAT5 signaling hallmark gene set, the significantly upregulated genes at 6h post activation and a list of RBP CLIP targets. **e)** CLIP data showing number and position of sequencing reads (in red) across *Cish* and *Socs1* transcripts (a set of top two lanes). In each set top lane shows ZFP36L1 CLIP data from OT-I CD8 CTLs stimulated for 3h with N4 peptide; bottom lane shows pan ZFP36 family CLIP data from in vitro activated naive CD4 T cells. ATTT and ATTTA motifs are identified in orange and blue. **f)** Schematic view of RBP target genes involved in IL2 signaling. Asterisks indicate direct RBP targets. Genes with icons highlighted in blue are differentially increased in dKO versus WT during the stimulation timecourse.

### Ablation of *Socs1* restores competitive fitness of RBP deficient T cells

The SOCS family proteins not only regulate cytokine responses, as some of its members such as *Cish* are also regulators of TCR signalling^41,44^. We deleted *Cish*, using Cas9 RNPs in naïve dKO OT-I cells (**Supplementary Figure 3c**) and tested whether absence of regulation of TCR signalling mediated by CISH would restore the dKO T cells competitiveness. However, co-culture of WT OT-I cells with *Cish* deficient dKO OT-I cells, did not rescue the dKO cells (**Fig.5a**). Next, we tested whether releasing the suppression of cytokine signaling in dKO cells by deletion of *Socs1* would improve the fitness of dKO cells. We deleted *Socs1* using Cas9 RNPs (**Supplementary Figure 3d**). Subsequent activation in co-culture with WT OT-I cells showed that dKO cells which received RNPs targeting *Socs1* expanded as well as WT OT-I cells in co-culture (**Fig.5b**). Expression of surface CD25 on dKO cells deficient for *Socs1*, as well as on co-cultured WT cells was significantly increased (**Fig.5c**). Thus, deletion of *Socs1* in RBP deficient T cells improves and prolongs their response to IL2 in co-culture. This specific de-repression of γ-cytokine signaling is sufficient to render the dKO cells competitive.

**Figure 5:**
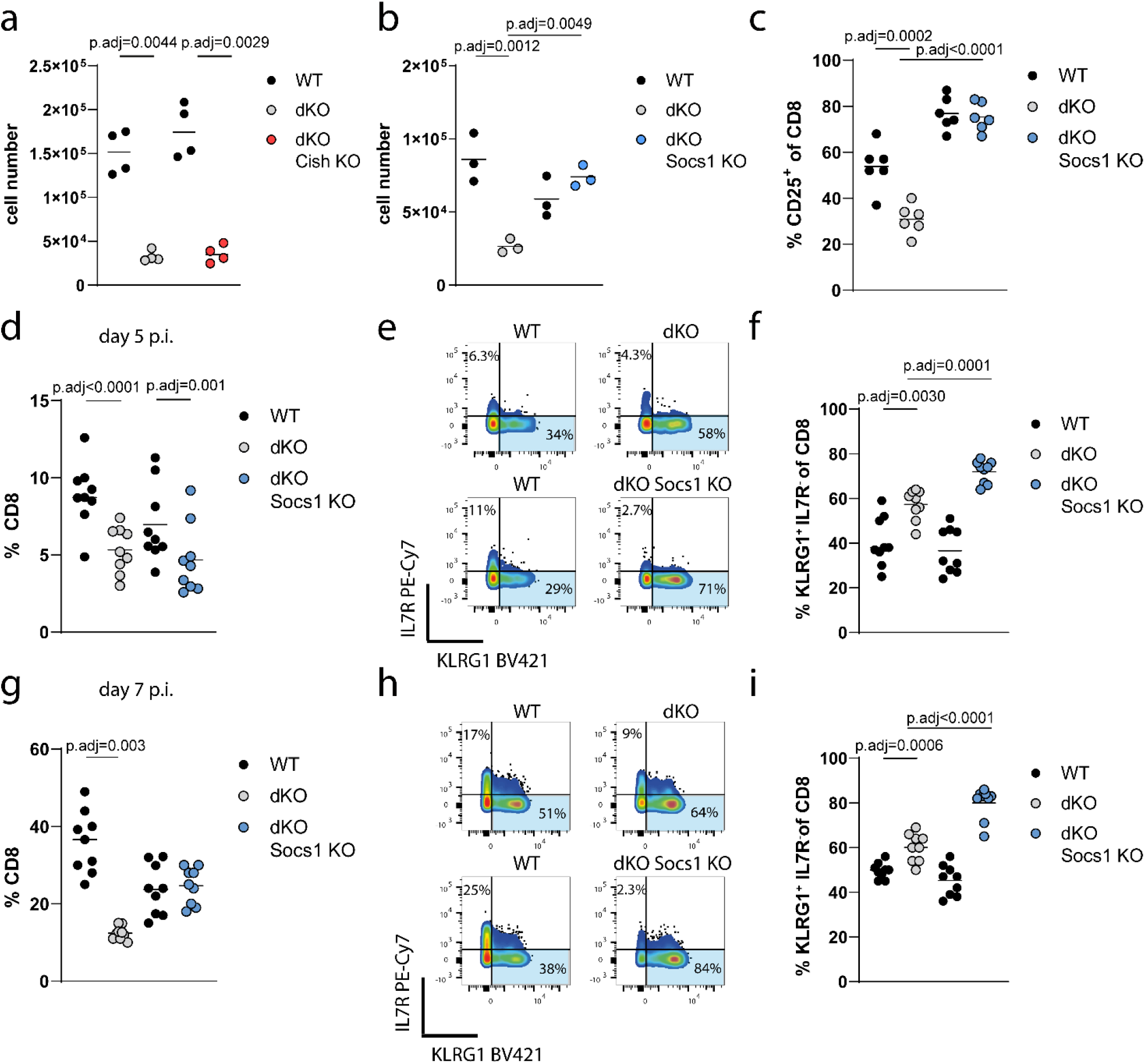
Ablation of *Socs1* restores competitive fitness of RBP deficient T cells. **a)** Cell numbers of WT OTI co-cultured with dKO OT-I cells or dKO OT-I cells where *Cish* or *Socs1* gene **b)** has been deleted 72h after stimulation with N4 peptide. Each dot in panels represents a technical replicate. Representative data from three independent experiments is shown. **c)** Frequencies of CD25 positive CD8 T cells are shown. Each dot in the panel represents a technical replicate and data is pooled from two of three independent experiments. **d)** Frequencies of total CD8 cells from co-transferred WT OT-I with either dKO OT-I or dKO OT-I cells where *Socs1* has been deleted in blood of recipient mice on day 5 after infection with *attLm*-OVA. **e)** Representative FACS plots and frequencies **f)** of KLRG1^+^ cells in blood of mice 5 days after infection. **g)** Frequencies of total CD8 cells from co-transfers in spleens of recipient mice on day 7 after infection with *attLm*-OVA. **h)** Representative FACS plots and frequencies **i)** of KLRG1^+^ cells in spleens/blood of recipient mice 7 days after infection. Statistical significance was determined by one-way ANOVA followed by Tukeys test for multiple comparisons. The statistical mean is indicated by horizontal line in all panels.

Next, we tested whether the release of cytokine signaling blockade by the deletion of *Socs1* in RBP deficient T cells restores their ability to compete with WT T cells *in vivo*. After co-transfer of WT and dKO OT-I cells treated with non-targeting guides or WT and dKO OT-I cells in which *Socs1* was deleted, we infected the recipient mice with *attLm*-Ova and followed the expansion of OT-I cells on day 5 in blood and day 7 in the spleen. On day 5 we found no clear effect of *Socs1* deletion on the expansion of dKO cells, which were still inferior to their WT counterparts (**Fig.5d**). Cells which were lacking *Socs1* had accumulated greater frequencies of KLRG1^+^ short lived effector cells (SLEC) (**Fig.5e, f**). Formation of SLEC is enhanced by the IL2-CD25 signaling axis^16,45^, suggesting that in the absence of *Socs1* the greater formation of SLEC is secondary to enhanced cytokine signalling. By day seven post infection, at the peak of the response, dKO OT-I cells lacking *Socs1* had expanded to the same extent as their WT counterparts, thus they no longer displayed a competitive disadvantage (**Fig.5g**). Also, on day 7 cells lacking *Socs1* had greater frequencies of KLRG1^+^ SLEC (**Fig.5h, i**). Overall the deletion of the RBP target *Socs1* in RBP deficient cells restores their competitive fitness allowing these cells to compete for IL2 and potentially other cytokines such as IL12^46^ and IL15^47^ *in vivo*. In addition, deletion of *Socs1* shows an additive effect on acceleration of SLEC formation by dKO cells. This exacerbation may result from the action of cytokine sensitive transcription factors which we previously identified as RBP targets which drive effector cell differentiation^30^.

### *ZFP36L1* plays a non-redundant role in mediating high affinity T cell expansion

To investigate how ZFP36 and ZFP36L1 contribute to T cell competition, we first assessed their expression in response to stimulation with peptides of varying affinity for the OT-I TCR. To trace the expression of both RBPs at single cell resolution we generated reporter mice in which open reading frames encoding the fluorescent proteins mAmetrine or mCherry were inserted in frame with the start codon of *Zfp36* and *Zfp36l1* respectively (**Supplementary Figure 4 a, b**). In these reporter mice, *mAmetrineZfp36* and *mCherryZfp36l1* are under the same transcriptional control as endogenous *Zfp36* and *Zfp36l1*. We validated that expression of ^mAmetrine^ZFP36 and ^mCherry^ZFP36L1 followed the same transient expression kinetics as the endogenous proteins using *in vitro* expanded and restimulated CD8 cells from OT-I ^mAmetrine/+^*Zfp36*^mCherry/+^*Zfp36l1* heterozygous mice (**Supplementary Figure 4 c, d**).

We stimulated naïve CD8 T cells from OT-I ^mAmetrine/+^*Zfp36*^mCherry/+^*Zfp36l1* heterozygous mice with variants of SIINFEKL peptide, which have altered stimulation potency of the OT-I TCR but bind equally to H-2K^b^. For stimulation we used splenocytes pulsed with different concentrations of high affinity N4 peptide and its low affinity variants Q4 (SIIQFEKL), T4 (SIITFEKL), which marks the threshold of positive and negative selection, and V4 (SIIVFEKL) which has a 1000 times lower potency compared to N4^48,49^. We measured the expression of ^mAmetrine^ZFP36 and ^mCherry^ZFP36L1 by Flow Cytometry. Frequencies of activated cells positive for ^mAmetrine^ZFP36 reached maximal levels with 0.3nM of N4,Q4 and T4, but required 3nM V4 (**Fig.6a**). In contrast, higher amounts of the lowest affinity peptide, (10nM V4), failed to induce a maximal frequency of ^mCherry^ZFP36L1 expressing cells (**Fig.6a**). Strikingly, the different behaviour in expression was especially evident when the mean fluorescence intensity per cell of each marker was analysed. High amounts of N4, Q4 and T4 peptides induced maximal expression of ^mAmetrine^ZFP36 per cell, as the fluorescence intensity reached a plateau irrespective of peptide affinity (**Fig.6b**). By contrast, ^mCherry^ZFP36L1 expression never reached the maximal levels observed after N4 stimulation even at high concentrations of Q4, T4 and V4 (**Fig.6b**). ^mCherry^ZFP36L1 expression per cell after stimulation with 10nM peptide was linearly related to peptide affinity and showed discriminative sensitivity for peptide variants (**Fig.6c, d**). This prompted us to test if both ZFP36 and ZFP36l1 are required for T cell competitiveness in response to high affinity antigen during infection. Therefore, we co-transferred naive OT-I specific WT cells with OT-I cells which lacked either ZFP36 or ZFP36L1 and infected the recipient mice with *attLm*-OVA. While CD45.1 WT and CD45.2 WT or ZFP36 KO OT-I cells expanded to same extent by day seven in the spleen of recipient mice, cells lacking only ZFP36L1 recapitulated the phenotype of the dKO cells in competition with WT OT-I cells (**Fig.6e**).

**Figure 6:**
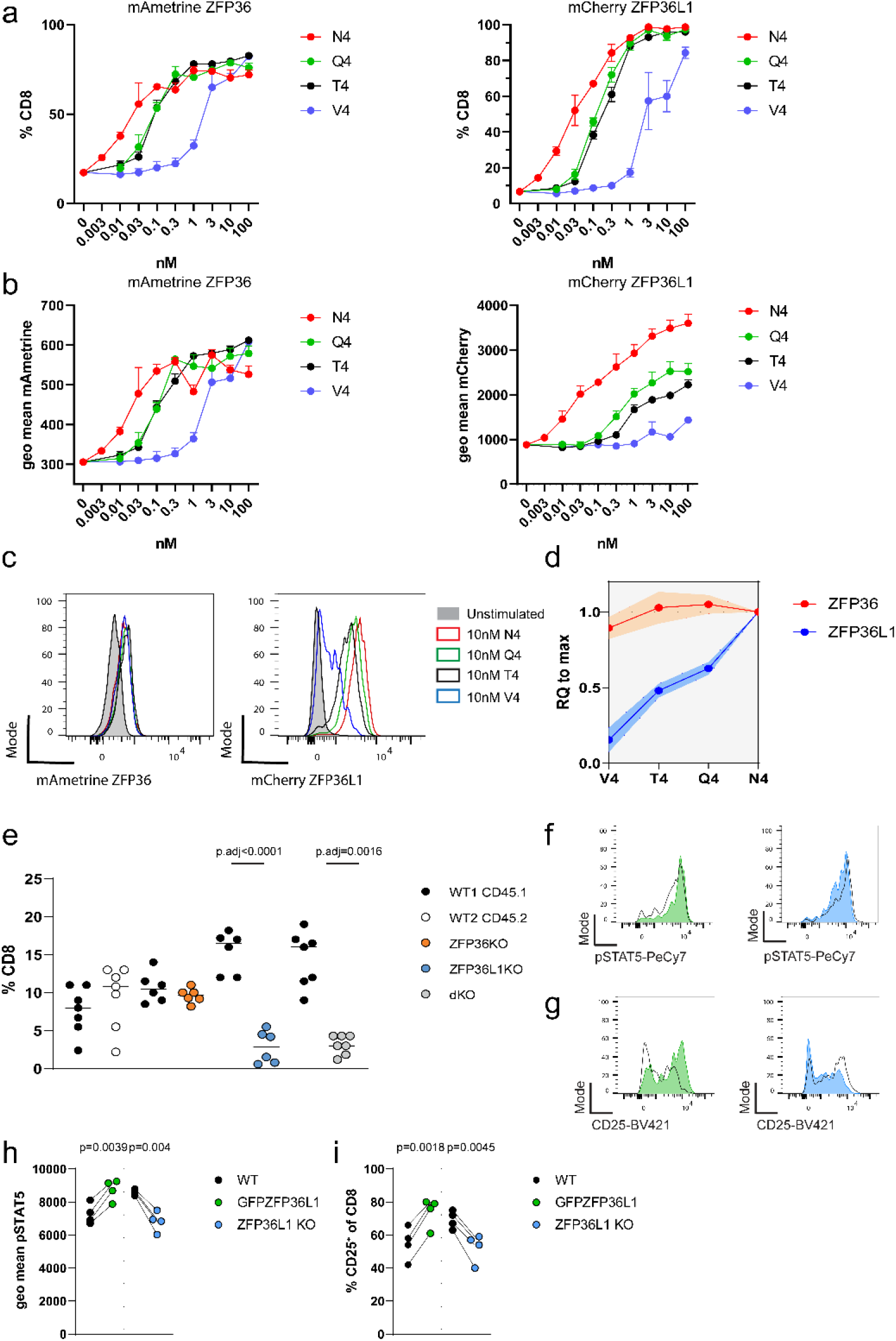
*ZFP36L1* plays a non-redundant role in mediating high affinity T cell expansion. **a)** Frequencies of ^mAmetrine^ZFP36^+^ (left panel) and ^mCherry^ZFP36L1^+^(right panel) CD8 T cells 6 hours after stimulation with peptides at indicated concentrations. **b)** Average expression per cell shown as geo mean of the expression of ^mAmetrine^ZFP36 (left panel) and ^mCherry^ZFP36L1 (right panel), when gated on the positive population. **c)** Representative histograms show expression ^mAmetrine^ZFP36 (left panel) and ^mCherry^ZFP36L1 (right panel), when stimulated with 10nM N4, V4 or T4 peptide. **d)** Relative expression (RQ) of ^**m**Ametrine^ZFP36 and ^mCherry^ZFP36L1. RQ was calculated as average mean expression of the fluorescent proteins per cell (geo mean) divided by their average expression when cells were stimulated with 10nM N4 peptide, which represents their maximal expression. Data in a), b) is compiled from three biological replicates and is representative of 3 independent experiments two of which included stimulation with Q4 peptide. Error bars indicate the standard deviation from the mean. Data in **d)** is compiled of the data from three independent experiments. Shaded area indicates the standard deviation from the mean. **e)** Frequencies of co-transferred OT-I cell in spleens of recipient mice, seven days after infection with *attLm*-OVA. Data is compiled from two independent experiments. Statistical significance was determined by two-way ANOVA followed by Tukey’s test for multiple comparisons. **f)** Representative histograms of pSTAT5 and **g)** CD25 in WT cells co-cultured with ZFP36L1 KO or GFPZFP36L1 cells 48 hours after activation with plate bound CD3 and CD28. **h)** Quantification of pSTAT5 **i)** CD25 expression. Each data point represents a technical replicate and data is representative for two independent experiments. Statistical significance was tested by paired t-test. All error bars indicate the standard deviation from the mean. The statistical mean is indicated by horizontal line in all panels.

To test further the role of ZFP36L1 in promoting STAT5 signaling we co-cultured naïve WT CD8 T cells with naïve CD8 T cells from transgenic mice which express a GFPZFP36L1 fusion protein^50^ (*GFPZfp36l1*). This model of forced expression of ZFP36L1, which is independent of TCR signalling, in addition to the endogenous expression of ZFP36L1 in these T cells, can be regarded as an overexpression system which we predicted to show the opposite effects on STAT5 signaling to the knockout. After activation with plate bound anti-CD3 and anti-CD28 for 48 hours we found pSTAT5 as well as CD25 expression in GFPZFP36L1 cells was greater than that in WT cells (**Fig.6f-i**). The opposite result was observed in ZFP36L1 KO cells where the expression of pSTAT5 and CD25 was reduced when compared to WT (**Fig.6f-i**). Taken together, these data show that ZFP36L1 promotes signalling via STAT5.

### *Socs1* limits the competitiveness of low affinity T cells

We next addressed whether enhancing the response to cytokines during an infection would improve the competitive fitness of low affinity CD8 T cells. We used OT-3 CD8 T cells which express a transgenic TCR with a 100-fold lower affinity for SIINFEKL than OT-I^51^. Co-transfer of OT-I and OT-3 cells followed by infection with *attLm*-OVA resulted in a 3x difference of transferred cell frequencies in the blood of recipient mice on day five post infection (**Fig.7a**). This difference was only slightly less upon co-transfer OT-I cells with OT-3 cells with deleted *Socs1* (**Fig.7a**). The deletion of *Socs1* in OT-3 cells led to an increased accumulation of KLRG1^+^ SLEC on day five post infection (**Fig.7b, c**). At the peak of the primary response, the disadvantage of OT-3 was more than a hundred-fold difference (**Fig.7d**). However, when OT-3 cells lacked *Socs1* this difference was significantly reduced (15 fold) (**Fig.7d**), suggesting that deletion of *Socs1* increases competitive fitness of low affinity CD8 T cells during the later expansion phase of the primary response. Thus, this result resembled the results observed with Socs1 deletion in dKO OT-I cells. On day seven, low affinity T cells lacking *Socs1* showed a greater tendency to form SLEC which is in contrast to WT OT3 cells which showed only a mild increase in KLRG1^+^ cells compared to WT OT-I (**Fig.7e, f**). This suggests that *Socs*1 limits expansion and differentiation into SLEC, which can be attributed to attenuation of cytokine signalling. In summary, we suggest that lifting the limits of cytokine signalling also enhances the competitiveness of low affinity CD8 T cells and therefore differential expression of ZFP36L1 should also contribute to their selection.

## Discussion

This work establishes ZFP36L1 as a mechanistic link between TCR affinity and the responsiveness to IL2. Moreover, this work explains previously observed priming defects of T cells and their failure to be efficiently recruited into immune responses in the absence of multiple ZFP36 family members^29,30^. Here, we show a T cell intrinsic mechanism which reveals ZFP36L1 to be a non-redundant regulator of cytokine signalling in T cells and key driver of cytokine driven clonal expansion.

The ability of the RBP to establish the competitive advantage of high affinity T cells is most apparent when RBP deficient high affinity OT-I cells have to compete with an equal number of high affinity WT OT-I T cells. The competitive disadvantage of cells lacking RBPs is much less pronounced when high affinity dKO OT-I T cells have to compete with endogenous T cells, with varying affinities for OVA derived antigens. The number of endogenous SIINFEKL K^b^ specific T cells is estimated to be no more than a few hundred^52^ and is composed of a range of affinities for SIINFEKL peptide. In line with this we find it striking that the expression of ZFP36L1 is highly responsive to the affinity of TCR peptide MHC-I interaction. We observed that antigen dose did not saturate expression of ZFP36L1 per cell. The high sensitivity of ZFP36L1 expression to changes in affinity suggests that it can act as a rheostat guiding the selection of high affinity clones by promoting their responsiveness to IL2. Similar TCR responsive expression behaviour has been reported for *Irf4* which converts TCR signals in an analogue fashion and drives metabolic reprogramming and expansion of CD8 T cells^12,53^. This contrasts with Myc which shows invariable expression per cell in response to TCR stimulation strength^11^ and Nur77 which marks an invariable activation threshold for T cells^10^.

ZFP36 and ZFP36L1 limit the expression of cytokines including IL2^30^, while ZFP36L1 specifically acts on the JAK-STAT5 pathway downstream of common γ-chain cytokine receptors. In this way ZFP36L1 is a crucial part of a type 2-incoherent feed forward loop^54^ controlling the IL2 signaling axis (**Fig. 7g**). In wild type cells the IL2 regulatory circuit would critically depend on exogenous input of IL2 to shift the equilibrium to enhance IL2 signaling. ZFP36L1 suppresses IL2 production while at the same time increasing sensitivity to it by suppressing a set of inhibitors of cytokine signalling. In the absence of ZFP36L1 sensitivity to IL2 is lost progressively, ultimately resulting in premature downregulation of CD25 and termination of a potent IL2 signal. This would be manifested most strikingly when IL2 becomes limiting during the course of an immune response^45^ and result in reduced clonal expansion, which is observed in the later expansion phase. ZFP36L1 could act in concert with the transcription factor IRF4 which shows similar TCR dependent regulation and promotes the cytokine response by inducing the expression of the β-chain of the IL2 receptor CD122^55^.

**Figure 7:**
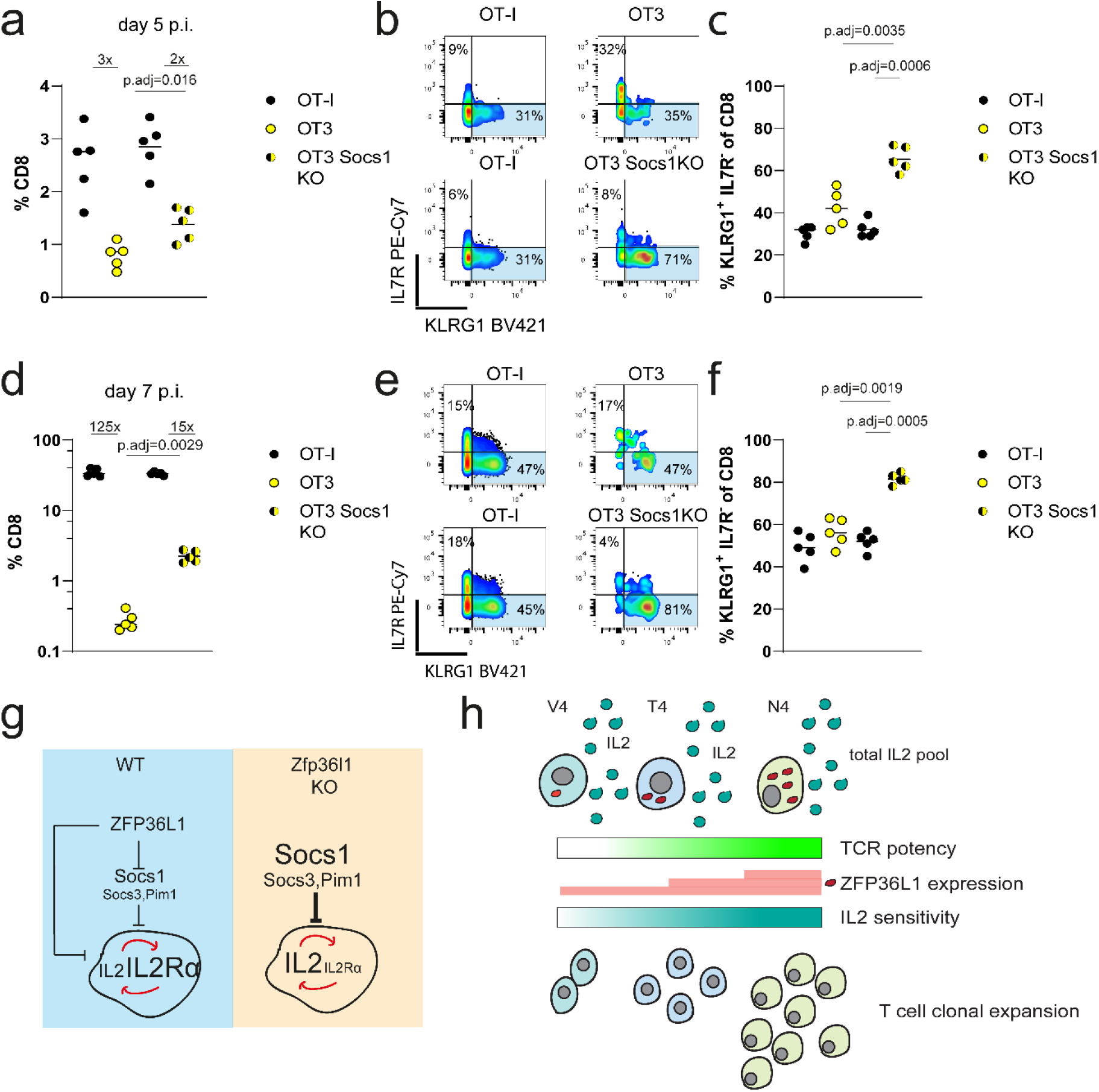
*Socs1* limits the competitiveness of low affinity T cells. **a)** Frequencies of total CD8 cells from co-transferred OT-I and OT-3 cells or OT-I and *Socs1* deficient OT-3 cells, in blood of recipient mice on day 5 after infection with *attLm*-OVA. **b)** Representative FACS plots and frequencies **c)** of KLRG1^+^ cells in blood of mice 5 days after infection. **d)** Frequencies of CD8 cells from co-transferred OT-I and OT-3 cells or OT-I and *Socs1* deficient OT-3 cells, in blood of recipient mice on day 7 after infection with *attLm*-OVA. **e)** Representative FACS plots and frequencies **f)** of KLRG1^+^ cells in blood of mice 7 days after infection. Fold differences between frequencies of cells are indicated in panels **a)** and **d)**. The statistical mean is indicated by horizontal line in all panels. Data is representative of two independent experiments. Statistical significance was determined by two-way ANOVA followed by Tukey’s test for multiple comparisons. Fold changes of the difference are indicated. **g)** Schematic model of ZFP36L1 comprising a crucial part of the IL2 signaling pathway as part of an incoherent feed forward regulatory loop. **h)** Schematic illustration of ZFP36L1 governing the selection and outgrowth of high affinity clones based on their sensitivity to IL2.

It has previously been shown that the affinity of the TCR does not necessarily drive the rate of initial cell division but that high affinity T cells proliferate and expand for an extended period of time^15^. Extended expansion of T cells is driven by cytokines including IL2. This becomes rate limiting during the course of an infection and efficient competition for IL2 requires the persistent expression of CD25^16^ which can be enhanced by other cytokines such as type-I interferons and IL12^8^ and promotes additional rounds of cell division. The ability of inflammatory cytokines to rescue high but not low affinity clones from apoptosis has been suggested as a mechanism for preferential outgrowth of high affinity clones when antigen availability is limited^17^. We show that ablation of *Socs1* not only restored the competitiveness of dKO cells but also that of T cells bearing a low affinity TCR in a competition with high affinity clones. This highlights that efficient sensing of IL2, and potentially other inflammatory signals, including IL12 and Interferon, is principally acting during competition and selection of high and low affinity T cells.

The limiting of autocrine IL2 production and the simultaneous de-repression of IL2 signaling suggests that ZFP36L1 enforces dependence of CD8 T cells on paracrine IL2, which becomes available during the early phase of an infection and is largely produced by antigen specific CD4 T cells and DCs^56^. We suggest that expression of ZFP36L1 in activated T cells significantly decreases autonomy during priming, increasing the dependence on costimulation but renders them competitive with CD4 effectors or regulatory T cells for available IL2 pools. In a situation where stimulation of T cells of varying affinities by antigen is sufficient to induce activation and proliferation, lesser expression of ZFP36L1 will position lower affinity T cells at a disadvantage to compete for cytokines, while at the same time safeguarding the costimulatory dependency and thus control/deletion of autoreactive and harmful clones (**Fig.7h**). Even small differences in IL2 sensitivity would result in the progressive loss of inferior clones; these would be outcompeted by more sensitive clones which are then further rewarded by having expanded more on the principle of competitive exclusion.

## Materials and Methods

### Mice

Mice with single or combined floxed *Zfp36* and *Zfp36l1* alleles^57,58^, B6.Cg-Tg(CD4-cre)1Cwi^59^ mice, GFP*Zfp36l1*^50^ and OT-I^60^ TCR (Vα2 and Vβ5 recognizing peptide residues 257-264 of chicken ovalbumin in the context of H2K^b^) were generated on the C57BL/6 background at the Babraham Institute. The B6.SJL-*Ptprc*^*a*^*Pepc*^*b*^/BoyJ (CD45.1) mice were bred at the Babraham Institute. CD45.1 CD45.2 double positive mice were bred as F1 from C57BL/6 (CD45.2) and B6.SJL-*Ptprc*^*a*^*Pepc*^*b*^/BoyJ (CD45.1) mice at the Babraham Institute. ^*mAmetrine*^*Zfp36* and ^*mCherry*^*Zfp36l1* mice were generated by Cyagen bioscience on a C57BL/6 background and maintained in the Babraham Institute Biological Support Unit.

Targeting vectors were designed to harbor the sequence encoding mAmetrine or mCherry without stop codon, in-frame upstream of the ATG translation start site of *Zfp36* or *Zfp36l1*, respectively. The *Zfp36l1* targeting vector featured diphtheria toxin A, a neomycin resistance cassette flanked by *Frt* recombination sequences and homology arms which were amplified from a mouse genomic BAC clone. The *ZFP36* targeting vector contained self-deletion anchor sites instead of *Frt* recombination sites. When homozygous, we did not observe lethality or any pathology in the reporter mice, suggesting a normal function of the proteins, deficiency of which causes an inflammatory syndrome^61^ or embryonic lethality^62^ in mice.

No primary pathogens or additional agents listed in the FELASA recommendations have been confirmed during health monitoring since 2009. Ambient temperature was ∼19–21°C and relative humidity 52%. Lighting was provided on a 12hr light: 12hr dark cycle including 15 min ‘dawn’ and ‘dusk’ periods of subdued lighting. After weaning, mice were transferred to individually ventilated cages with 1–5 mice per cage. Mice were fed CRM (P) VP diet (Special Diet Services) *ad libitum* and received seeds (e.g., sunflower, millet) at the time of cage-cleaning as part of their environmental enrichment. All mouse experimentation was approved by the Babraham Institute Animal Welfare and Ethical Review Body. Animal husbandry and experimentation complied with existing European Union and United Kingdom Home Office legislation.

### Infection with *Listeria monocytogenes*

8-14-week old male and female mice were used for all experiments. Bacteria were grown in BHI medium to an OD_600_ of 0.1 before each experiment. Mice were infected with a sublethal dose of 5×10^6^ CFU attenuated (Δ*actA*) *listeria monocytogenes* expressing OVA_257-264_ ^30^by intravenous administration.

Adoptive transfer experiments: naive OT-I cells (if not otherwise stated) were sorted by Flow Cytometry from spleens and LN of mice from a respective genotype and a total of 1000 cells per mouse transferred intravenously on the day before infection. In all cotransfer experiments donor cells were mixed with 1*10^6^ carrier splenocytes of the same genotype as the host.

For characterization of transferred OT-I cells after transfer mice were bled on indicated dates and spleens were collected at final points of analysis. For analysis of cytokine production, cells were incubated with 10^−7^M N4 peptide for 3 hours in the presence of Brefeldin A (1μg/ml) in full cell culture medium (IMDM including 10%FCS 50μM beta β-Me). After surface staining, cells were fixed with 2% PFA for 20 min at 4°C and permeabilized with BD Perm/wash +1%FCS for 20 min at 4°C, before intracellular Cytokine staining.

### *In vitro* culture and activation of T cells

All *in vitro* and cell culture experiments were performed in IMDM culture medium (Gibco) supplemented with 10% FCS, GlutaMAX^™^ (Gibco), 40U Penicillin /Streptomycin (Gibco) and 50μM β-Mercaptoethanol.

Naive CD8 T cells were isolated from spleen and LN using two rounds of negative depletion with Streptavidin DynaBeads^™^ (Thermo Fisher) using 1*10^8^ and 4*10^7^ beads per 1*10^8^ total splenocytes/LN cells. The following antibodies were used for the depletion cocktail: B220-Bio (RA3-6B3), CD4-Bio (GK1.5), CD11b-Bio (M1/70), CD11c-Bio (N418), CD19-Bio (1D3), CD44-Bio (IM7), CD105-Bio, F4/80-Bio (BM8), GR1-Bio (RB6-8C5), NK1.1-Bio (PK136), Ter119-Bio (Ter-119), γδTCR-Bio (GL3).

Cells were stimulated at a starting concentration of 5*10^5^ cells/ml, if not otherwise stated with 5μg/ml plate bound anti-CD3 (145-2C11, BioXcell) and 1μ/ml plate bound anti-CD28 (37.51, BioXcell). OT-I transgenic naive CD8 T cells were stimulated with (N4) SIINFEKL peptide at 10^−10^M if not otherwise stated. The altered peptide ligands SIIQFEKL, SIITFEKL and SIIVFEKL were used to stimulate OT-I cells in some experiments. In some conditions recombinant mouse IL2 and IL12 (Peprotech) were added at 20ng/ml and 2.5ng/ml respectively or at concentrations as indicated. Anti-mouse IL2 (JES6-1A12, BioXcell) was used at 50ug/ml to block autocrine IL2. Recombinant human IL2 (Peprotech) was added at indicated concentrations.

For the analysis of proliferation, cells were labelled with cell trace violet (Thermo Fisher) or cell trace yellow (Thermo Fisher) at 10μM final concentration for 6 min at 37°C, before stimulation. Cell numbers per generation were enumerated using cell counting beads for flow cytometry (ACBP-50-10, Spherotech).

### CrispR Cas9 RNP mediated gene editing

For electroporation of Cas9 Ribonucleoproteins naïve OT-I T cells were isolated using the StemCell EasySep Mouse Naïve CD8^+^ T Cell Isolation Kit (StemCell; 19853). Isolated cells were electroporated with complexes of Cas9 and control or IL2 targeting gRNA (all from IDT) in OptiMEM medium (Gibco) using the NEPA21 electroporator (Nepagene). After electroporation cells were cultured for 24h in RPMI medium (Gibco) containing 10% FCS and 10ng/ml recombinant IL7 (Peprotech; 217-17), before transfer into recipient mice. The following guide sequence (IDT) was used to target the indicated genomic sequence of the *Socs1* gene: AAGTGCACGCGGATGCTCGT, *Il2* gene: AAGATGAACTTGGACCTCTG *Cish* gene: CTTGTCAAGACCTCGAATCC.

### Flow Cytometry and antibodies

For cell surface staining single cell suspensions from tissues or cultured cells were prepared in FACS buffer containing 1x PBS, 1% FCS ± 2 mM EDTA (if not otherwise stated in the methods). All cells were blocked with Fcγ blocking antibody (24G2, BioXcell) and incubated with fixable cell viability dye eF780 (Thermo Fisher or BD) for 20 min at 4 °C. For intracellular staining cells were fixed with BD Cytofix/Cytoperm (554722) or 2–4% PFA for 20 min at 4 °C. Cells were permeabilized with BD Permwash (554723) containing 1% FCS for 20 min at 4 °C. Intracellular staining was performed in BD Permwash containing 0.5% FCS and the intracellular antibody cocktail for 1h at RT. Surface stained cells from infection experiments were fixed with BD Cytofix/Cytoperm for 30 min at 4 °C before analysis. Staining for pSTAT5 and anti-apoptotic molecules was performed by fixing cells immediately during stimulation with 4% PFA at final concentration of 2% on ice. Cells were fixed for 30min. Cells were permeabilized with 90% Methanol for 30 min on ice and stored at −20 °C. Cells were finally stained in 1xPBS + 0.05% BSA for 1h on ice.

The following antibody clones were used in the flow cytometry experiments: CD8 (53-6.7;1:400), CD45.1 (A20;1:100), CD45.2 (104;1:100), KLRG1 (2F1;1:400), CD127 (A7R34;1:100), CD44 (IM7;1:400), CD25 (7D4; PC61;1:400), CD69 (H1.2F3;1:400), TCRβ (H57-597;1:200), Ki-67 (SolA15;1:100), Mcl1 (LUVBKM; 1:20), Bcl2 (BCL/10C4; 1:50), pSTAT5 (47;1:5)), activeCaspase3 (C92-605;1:100). CD16/32 2.4G2 BioXcell BE0008 (1:2000, 0.5μg/ml). Data was acquired using a Fortessa Flow Cytometer equipped with 355 nm, 405 nm, 488 nm, 561 nm and 640 nm lasers (Beckton Dickinson). Flow Cytometry data was analysed using FlowJo 10.6 software.

### Western Blotting

Whole cell lysates were prepared by resuspending CTL cell pellets in 2x Laemmli buffer containing 5% β2-mercaptoethanol. The equivalent of 1-2 million cells were loaded per lane. Protein concentrations were determined by BCA protein assay (Pierce,23225). Samples were resolved by 12 % SDS-PAGE and transferred to nitrocellulose membrane using iBlot2 transfer device (IB21001). The following antibodies were used in western blotting experiments: Rabbit anti-CISH (D4D9; 1:100) (Cell signalling), Goat polyclonal anti-SOCS1 (ab9870; 1:500) (Abcam), mouse anti Tubulin (DM1A;1:10000) (Sigma). Secondary HRP conjugated antibodies: anti goat HRP Trueblot (1:2000) (ebioscience), anti-rabbit HRP (1:10000) (DAKO). Mouse anti ZFP36 (Origene #OTI3D10, 2μg/ml), rabbit anti ZFP36L1 (CST #BRF1/2, 32ng/ml), rabbit anti GAPDH (CST #5174, D16H11, 1:1000). Secondary antibodies for detection with Licor used: anti-Mouse IgG IRDye800CM (Licor #926-32210) and anti-Rabbit IgG IRDye680RD (Licor #925-68071). Membranes were scanned using Licor Odyssey CLx using standard methods, or ECL prime (Cytiva, RPN2232) for HRP conjugated antibody detection. Image analysis was conducted using ImageStudio Lite version 5.2, and normalized protein signal was calculated using standard methods.

### RNA sequencing

500.000 OT-I-transgenic naïve CD8+ T cells were stimulated in 200ul IMDM medium containing 10%FCS, 50μM beta-Me and 40U Penicillin /Streptomycin with (N4) SIINFEKL peptide at 10 ^−10^M, for 0-16h. Total RNA was extracted using the RNeasy Micro Kit (Qiagen, REF:74004). Opposing strand-specific RNA-seq libraries were generated using the SMARTer Stranded Total RNA-Seq Kit v3 - Pico Input Mammalian (Takara), according to the manufacturer’s instructions, and sequenced on an Illumina NovaSeq 6000 using 150bp paired-end reads.

### Data and statistical analysis

Statistical analysis was performed using Graph Pad Prism 9.3.1 and Microsoft Excel. RNA-seq data from in-vitro activated CD8^+^ T cells was trimmed using Trim Galore (https://www.bioinformatics.babraham.ac.uk/projects/trim_galore), first to remove poor quality and adapter sequence from the 3’ end of all reads, and second to remove 14 bases from the 5’ end of read 2 and 11 bases from the 3’ end of read 1, to ensure removal of the in-line UMI sequence comprising the start of read 2. Reads were mapped to the GRCm39 mouse genome using HISAT2 (v2.1.0), suppressing unpaired or discordant alignments and considering known splice sites from the Ensembl GRCm39 v103 annotation. Note that this resulted in a much lower mapping efficiency and thus fewer mapped reads than expected; coupled with our quality control results and further mapping to rRNA sequences using Bowtie, substantial ribosomal RNA contamination was identified as the cause of this low mapping efficiency, suggesting inefficient depletion during library preparation. The quality and consistency between replicates for the successfully mapped reads, however, appeared good, therefore we continued with the analysis. Raw read counts over mRNA features were quantified using Seqmonk (https://www.bioinformatics.babraham.ac.uk/projects/seqmonk). Differentially expressed genes at the different timepoints after CD8^+^ T cell activation were identified using DESeq2 analysis with default parameters, comparing dKO to WT for each time point, and significantly increased genes were selected for adjusted p value <= 0.05 and log2 FC > 0.3 (using ‘normal’ log2 fold change shrinkage). ZFP36/L1 library of CLIP targets for T cells was collated as previously described^30^. CLIP data was visualized using the following application: https://github.com/LouiseMatheson/iCLIP_visualisation_shiny.

## Supporting information

Supplementary Table 1 CLIP Target genes

## Acknowledgements

We thank Dr. Dietmar Zehn for providing the OT-3 transgenic mice. We also thank Dr. Arianne Richard, Dr. Peter Katsikis, Dr. Sarah Bell, Dr. Beatriz Saenz and Dr. Francesca Rossi for critically reading the manuscript. We would like to thank Sarah Collison for formatting this manuscript version for BioRxiv, using a template generously provided by the Finkelstein Lab, Austin, TX (https://github.com/finkelsteinlab). We thank the UKRI-BBSRC Core Capability Grant funded Babraham Institute Biological Support Unit, Sequencing, Flow Cytometry and Bioinformatics Facilities for support. This study was additionally funded by the BBSRC (BB/P01898X/1; BBS/E/B/000C0407; BBS/E/B/000C0428; the BBSRC to the Babraham Institute; and a Wellcome Investigator award (200823/Z/16/Z) to M.T.

## Author Contributions

G.P. conceptualization; methodology; investigation; validation; formal analysis; visualisation; writing original draft preparation;

T.J.M., M.J.E. and F.S. methodology, validation, review and editing

L.M. software; formal analysis; investigation; visualisation; review and editing;

M.T. Conceptualisation, supervision, funding acquisition, writing, review and editing.

## Data availability statement

Sequencing data of time course gene expression in WT and dKO CD8 T cells is publicly available on: GEO: GSE230373.

## References

1. Bennett, T. J., Udupa, V. A. V. & Turner, S. J. Running to Stand Still: Naive CD8+ T Cells Actively Maintain a Program of Quiescence. Int. J. Mol. Sci. 21, 9773 (2020).

2. Kavazović, I., Polić, B. & Wensveen, F. M. Cheating the Hunger Games; Mechanisms Controlling Clonal Diversity of CD8 Effector and Memory Populations. Front. Immunol. 9, 2831 (2018).

3. Wensveen, F. M., Gisbergen, K. P. J. M. & Eldering, E. The fourth dimension in immunological space: how the struggle for nutrients selects high-affinity lymphocytes. Immunol. Rev. 249, 84–103 (2012).

4. Zenke, S. et al. Quorum Regulation via Nested Antagonistic Feedback Circuits Mediated by the Receptors CD28 and CTLA-4 Confers Robustness to T Cell Population Dynamics. Immunity 52, 313–327.e7 (2020).

5. Thaventhiran, J. E. D. et al. Activation of the Hippo pathway by CTLA-4 regulates the expression of Blimp-1 in the CD8 + T cell. Proc. Natl. Acad. Sci. 109, (2012).

6. Uhl, L. F. K. & Gérard, A. Modes of Communication between T Cells and Relevance for Immune Responses. Int. J. Mol. Sci. 21, 2674 (2020).

7. Curtsinger, J. M. & Mescher, M. F. Inflammatory cytokines as a third signal for T cell activation. Curr. Opin. Immunol. 22, 333–340 (2010).

8. Starbeck-Miller, G. R., Xue, H.-H. & Harty, J. T. IL-12 and type I interferon prolong the division of activated CD8 T cells by maintaining high-affinity IL-2 signaling in vivo. J. Exp. Med. 211, 105–120 (2014).

9. Voisinne, G. et al. T Cells Integrate Local and Global Cues to Discriminate between Structurally Similar Antigens. Cell Rep. 11, 1208–1219 (2015).

10. Au-Yeung, B. B. et al. A sharp T-cell antigen receptor signaling threshold for T-cell proliferation. Proc. Natl. Acad. Sci. 111, (2014).

11. Preston, G. C. et al. Single cell tuning of Myc expression by antigen receptor signal strength and interleukin–2 in T lymphocytes. EMBO J. 34, 2008–2024 (2015).

12. Conley, J. M., Gallagher, M. P., Rao, A. & Berg, L. J. Activation of the Tec Kinase ITK Controls Graded IRF4 Expression in Response to Variations in TCR Signal Strength. J. Immunol. 205, 335–345 (2020).

13. Au-Yeung, B. B. et al. IL-2 Modulates the TCR Signaling Threshold for CD8 but Not CD4 T Cell Proliferation on a Single-Cell Level. J. Immunol. 198, 2445–2456 (2017).

14. Richard, A. C. et al. T cell cytolytic capacity is independent of initial stimulation strength. Nat. Immunol. 19, 849–858 (2018).

15. Zehn, D., Lee, S. Y. & Bevan, M. J. Complete but curtailed T-cell response to very low-affinity antigen. Nature 458, 211–214 (2009).

16. Obar, J. J. et al. CD4 + T cell regulation of CD25 expression controls development of short-lived effector CD8 + T cells in primary and secondary responses. Proc. Natl. Acad. Sci. 107, 193–198 (2010).

17. Zehn, D., Roepke, S., Weakly, K., Bevan, M. J. & Prlic, M. Inflammation and TCR Signal Strength Determine the Breadth of the T Cell Response in a Bim-Dependent Manner. J. Immunol. 192, 200–205 (2014).

18. Feau, S., Arens, R., Togher, S. & Schoenberger, S. P. Autocrine IL-2 is required for secondary population expansion of CD8+ memory T cells. Nat. Immunol. 12, 908–913 (2011).

19. Williams, M. A., Tyznik, A. J. & Bevan, M. J. Interleukin-2 signals during priming are required for secondary expansion of CD8+ memory T cells. Nature 441, 890–893 (2006).

20. D’Souza, W. N., Schluns, K. S., Masopust, D. & Lefrançois, L. Essential Role for IL-2 in the Regulation of Antiviral Extralymphoid CD8 T Cell Responses. J. Immunol. 168, 5566–5572 (2002).

21. Bachmann, M. F., Wolint, P., Walton, S., Schwarz, K. & Oxenius, A. Differential role of IL-2R signaling for CD8+ T cell responses in acute and chronic viral infections. Eur. J. Immunol. 37, 1502–1512 (2007).

22. D’Souza, W. N. & Lefrançois, L. IL-2 Is Not Required for the Initiation of CD8 T Cell Cycling but Sustains Expansion. J. Immunol. 171, 5727–5735 (2003).

23. Toumi, R. et al. Autocrine and paracrine IL-2 signals collaborate to regulate distinct phases of CD8 T cell memory. Cell Rep. 39, 110632 (2022).

24. Wensveen, F. M. et al. Apoptosis Threshold Set by Noxa and Mcl-1 after T Cell Activation Regulates Competitive Selection of High-Affinity Clones. Immunity 32, 754–765 (2010).

25. Höfer, T., Krichevsky, O. & Altan-Bonnet, G. Competition for IL-2 between Regulatory and Effector T Cells to Chisel Immune Responses. Front. Immunol. 3, (2012).

26. Busse, D. et al. Competing feedback loops shape IL-2 signaling between helper and regulatory T lymphocytes in cellular microenvironments. Proc. Natl. Acad. Sci. 107, 3058–3063 (2010).

27. Amado, I. F. et al. IL-2 coordinates IL-2–producing and regulatory T cell interplay. J. Exp. Med. 210, 2707–2720 (2013).

28. Salerno, F. et al. Translational repression of pre-formed cytokine-encoding mRNA prevents chronic activation of memory T cells. Nat. Immunol. 19, 828–837 (2018).

29. Cook, M. E. et al. The ZFP36 family of RNA binding proteins regulates homeostatic and autoreactive T cell responses. Sci. Immunol. 7, eabo0981 (2022).

30. Petkau, G. et al. The timing of differentiation and potency of CD8 effector function is set by RNA binding proteins. Nat. Commun. 13, 2274 (2022).

31. Matheson, L. S. et al. Multiomics analysis couples mRNA turnover and translational control of glutamine metabolism to the differentiation of the activated CD4+ T cell. Sci. Rep. 12, 19657 (2022).

32. Kim, H.-P., Kelly, J. & Leonard, W. J. The Basis for IL-2-Induced IL-2 Receptor α Chain Gene Regulation. Immunity 15, 159–172 (2001).

33. Moore, M. J. et al. ZFP36 RNA-binding proteins restrain T cell activation and anti-viral immunity. eLife 7, e33057 (2018).

34. Palmer, D. C. & Restifo, N. P. Suppressors of cytokine signaling (SOCS) in T cell differentiation, maturation, and function. Trends Immunol. 30, 592–602 (2009).

35. Davey, G. M., Heath, W. R. & Starr, R. SOCS1: a potent and multifaceted regulator of cytokines and cell-mediated inflammation. Tissue Antigens 67, 1–9 (2006).

36. Sporri, B., Kovanen, P. E., Sasaki, A., Yoshimura, A. & Leonard, W. J. JAB/SOCS1/SSI-1 is an interleukin-2–induced inhibitor of IL-2 signaling. Blood 97, 221–226 (2001).

37. Cohney, S. J. et al. SOCS-3 Is Tyrosine Phosphorylated in Response to Interleukin-2 and Suppresses STAT5 Phosphorylation and Lymphocyte Proliferation. Mol. Cell. Biol. 19, 4980–4988 (1999).

38. Tannahill, G. M. et al. SOCS2 Can Enhance Interleukin-2 (IL-2) and IL-3 Signaling by Accelerating SOCS3 Degradation. Mol. Cell. Biol. 25, 9115–9126 (2005).

39. Knosp, C. A. et al. SOCS2 regulates T helper type 2 differentiation and the generation of type 2 allergic responses. J. Exp. Med. 208, 1523–1531 (2011).

40. Landsman, T. & Waxman, D. J. Role of the Cytokine-induced SH2 Domain-containing Protein CIS in Growth Hormone Receptor Internalization. J. Biol. Chem. 280, 37471–37480 (2005).

41. Palmer, D. C. et al. Cish actively silences TCR signaling in CD8+ T cells to maintain tumor tolerance. J. Exp. Med. 212, 2095–2113 (2015).

42. Matsumoto, A. et al. Suppression of STAT5 Functions in Liver, Mammary Glands, and T Cells in Cytokine-Inducible SH2-Containing Protein 1 Transgenic Mice. Mol. Cell. Biol. 19, 6396–6407 (1999).

43. Peltola, K. J. et al. Pim-1 kinase inhibits STAT5-dependent transcription via its interactions with SOCS1 and SOCS3. Blood 103, 3744–3750 (2004).

44. Guittard, G. et al. The Cish SH2 domain is essential for PLC-γ1 regulation in TCR stimulated CD8+ T cells. Sci. Rep. 8, 5336 (2018).

45. Kalia, V. et al. Prolonged Interleukin-2Rα Expression on Virus-Specific CD8+ T Cells Favors Terminal-Effector Differentiation In Vivo. Immunity 32, 91–103 (2010).

46. Eyles, J. L., Metcalf, D., Grusby, M. J., Hilton, D. J. & Starr, R. Negative Regulation of Interleukin-12 Signaling by Suppressor of Cytokine Signaling-1. J. Biol. Chem. 277, 43735–43740 (2002).

47. Davey, G. M. et al. SOCS-1 regulates IL-15–driven homeostatic proliferation of antigen-naive CD8 T cells, limiting their autoimmune potential. J. Exp. Med. 202, 1099–1108 (2005).

48. Oberle, S. G. et al. A Minimum Epitope Overlap between Infections Strongly Narrows the Emerging T Cell Repertoire. Cell Rep. 17, 627–635 (2016).

49. Daniels, M. A. et al. Thymic selection threshold defined by compartmentalization of Ras/MAPK signalling. Nature 444, 724–729 (2006).

50. Vogel, K. U., Bell, L. S., Galloway, A., Ahlfors, H. & Turner, M. The RNA-Binding Proteins Zfp36l1 and Zfp36l2 Enforce the Thymic β-Selection Checkpoint by Limiting DNA Damage Response Signaling and Cell Cycle Progression. J. Immunol. 197, 2673–2685 (2016).

51. Enouz, S., Carrié, L., Merkler, D., Bevan, M. J. & Zehn, D. Autoreactive T cells bypass negative selection and respond to self-antigen stimulation during infection. J. Exp. Med. 209, 1769–1779 (2012).

52. Obar, J. J., Khanna, K. M. & Lefrançois, L. Endogenous Naive CD8+ T Cell Precursor Frequency Regulates Primary and Memory Responses to Infection. Immunity 28, 859–869 (2008).

53. Nayar, R. et al. Graded Levels of IRF4 Regulate CD8 + T Cell Differentiation and Expansion, but Not Attrition, in Response to Acute Virus Infection. J. Immunol. 192, 5881–5893 (2014).

54. Alon, U. Network motifs: theory and experimental approaches. Nat. Rev. Genet. 8, 450–461 (2007).

55. Huang, S. et al. IRF4 Modulates CD8+ T Cell Sensitivity to IL-2 Family Cytokines. ImmunoHorizons 1, 92–100 (2017).

56. Kalia, V. & Sarkar, S. Regulation of Effector and Memory CD8 T Cell Differentiation by IL-2—A Balancing Act. Front. Immunol. 9, 2987 (2018).

57. Hodson, D. J. et al. Deletion of the RNA-binding proteins ZFP36L1 and ZFP36L2 leads to perturbed thymic development and T lymphoblastic leukemia. Nat. Immunol. 11, 717–724 (2010).

58. Newman, R. et al. Maintenance of the marginal-zone B cell compartment specifically requires the RNA-binding protein ZFP36L1. Nat. Immunol. 18, 683–693 (2017).

59. Lee, P. P. et al. A Critical Role for Dnmt1 and DNA Methylation in T Cell Development, Function, and Survival. Immunity 15, 763–774 (2001).

60. Hogquist, K. A. et al. T cell receptor antagonist peptides induce positive selection. Cell 76, 17–27 (1994).

61. Taylor, G. A. et al. A Pathogenetic Role for TNFα in the Syndrome of Cachexia, Arthritis, and Autoimmunity Resulting from Tristetraprolin (TTP) Deficiency. Immunity 4, 445–454 (1996).

62. Bell, S. E. et al. The RNA binding protein Zfp36l1 is required for normal vascularisation and post-transcriptionally regulates VEGF expression. Dev. Dyn. 235, 3144–3155 (2006).

