## Supplementary Figures for "*Zfp36l1* establishes the high affinity CD8 T cell response by directly linking TCR affinity to cytokine sensing"

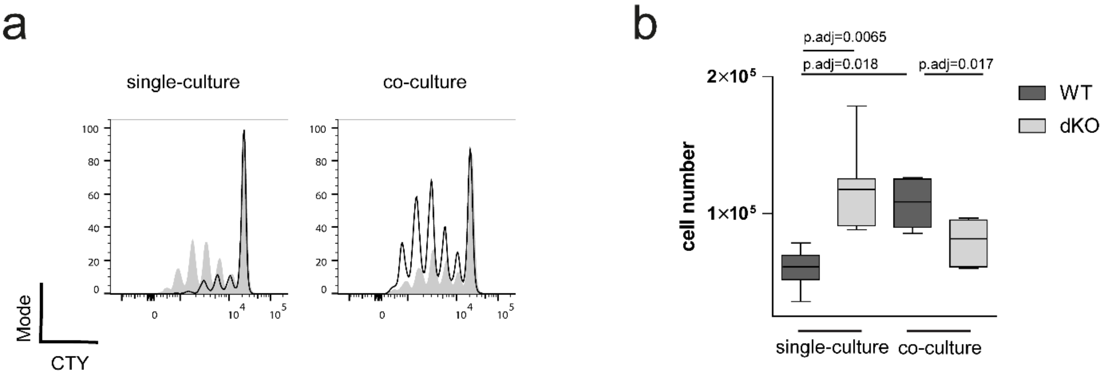


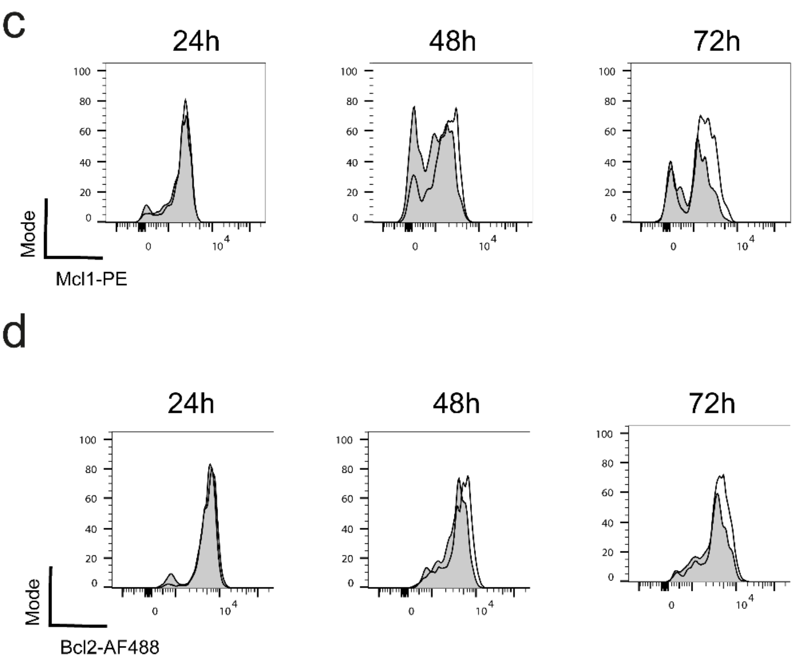


**Supplementary Figure 1: RBP deficient T cells promote the WT T cell response in co-culture**. **a)** Representative panels show the dilution of CTY by naïve WT (open) and dKO (filled) CD8 T cells stimulated with plate bound anti CD3 and anti CD28 for 72h individually or in co-culture **b)** Absolute cell numbers of WT and dKO cells after 72 hours with plate bound anti CD3 and anti CD28 in individual and co-culture. Data is compiled from 7 independent experiments. **c)** Representative histograms show Mcl1 and d) Bcl2 expression in WT (open histograms) and dKO (filled histograms) in co-cultures activated for indicated times with peptide. Statistical significance was determined by one-way ANOVA followed by Tukey`s test for multiple comparisons. Box Plots in **b)** indicate the data distribution by showing the min and max values around the statistical mean.

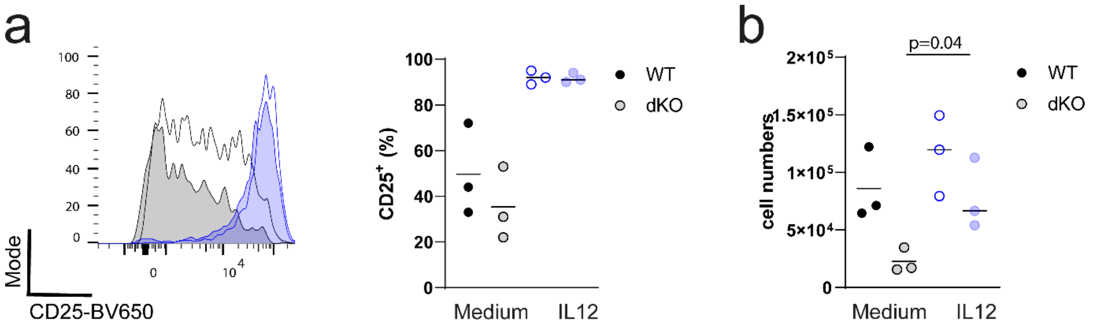


**Supplementary Figure 2: IL12 promotes dKO competitiveness and induces CD25 expression while autocrine IL2 production is not detrimental to T cell fitness**. **a)** Representative histograms show the expression of CD25 by WT (open) and dKO (filled) CD8 T cells stimulated with N4 in the presence (blue) or absence of IL12. **b)** Absolute cell numbers of WT and dKO OT-I cells after 72 hours stimulation with N4 peptide in co-culture in the presence or absence of IL12. Statistical significance was determined by unpaired t-test. The statistical mean is indicated by horizontal line in all panels.

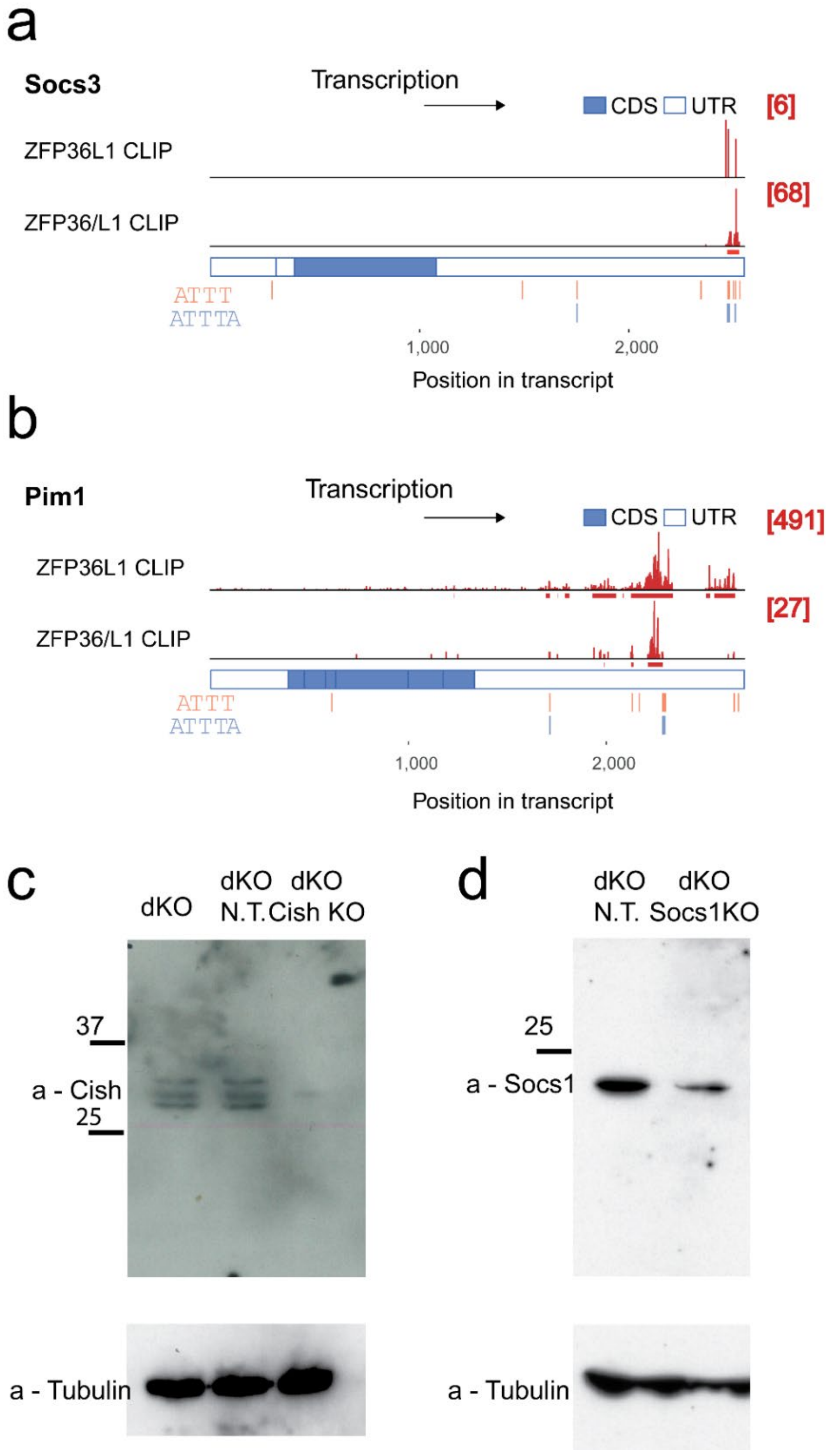


**Supplementary Figure 3: Socs3 and Pim1 are direct RBP CLIP targets and validation of Cas9 deletion of *Cish* and *Socs1*.** **a)** CLIP data showing number and position of sequencing reads (in red) across *Socs3* and *Pim1* **b)** transcripts (a set of top two lanes). In each set top lane shows ZFP36L1 CLIP data from OT-I CD8 CTLs stimulated for 3h with N4 peptide; bottom lane shows panZFP36 family CLIP data from in vitro activated naive CD4 T cells. ATTT and ATTTA motifs are identified in orange and blue. **c)** Validation of deletion of *Cish* and **d)** *Socs1* by western blot in OT-I CTLs differentiated for 6 days in the presence of IL2 after stimulation with N4 peptide. Lanes were loaded with lysates from dKO cells treated with non-targeting guides or specific guides for *Cish* and *Socs1* as indicated.

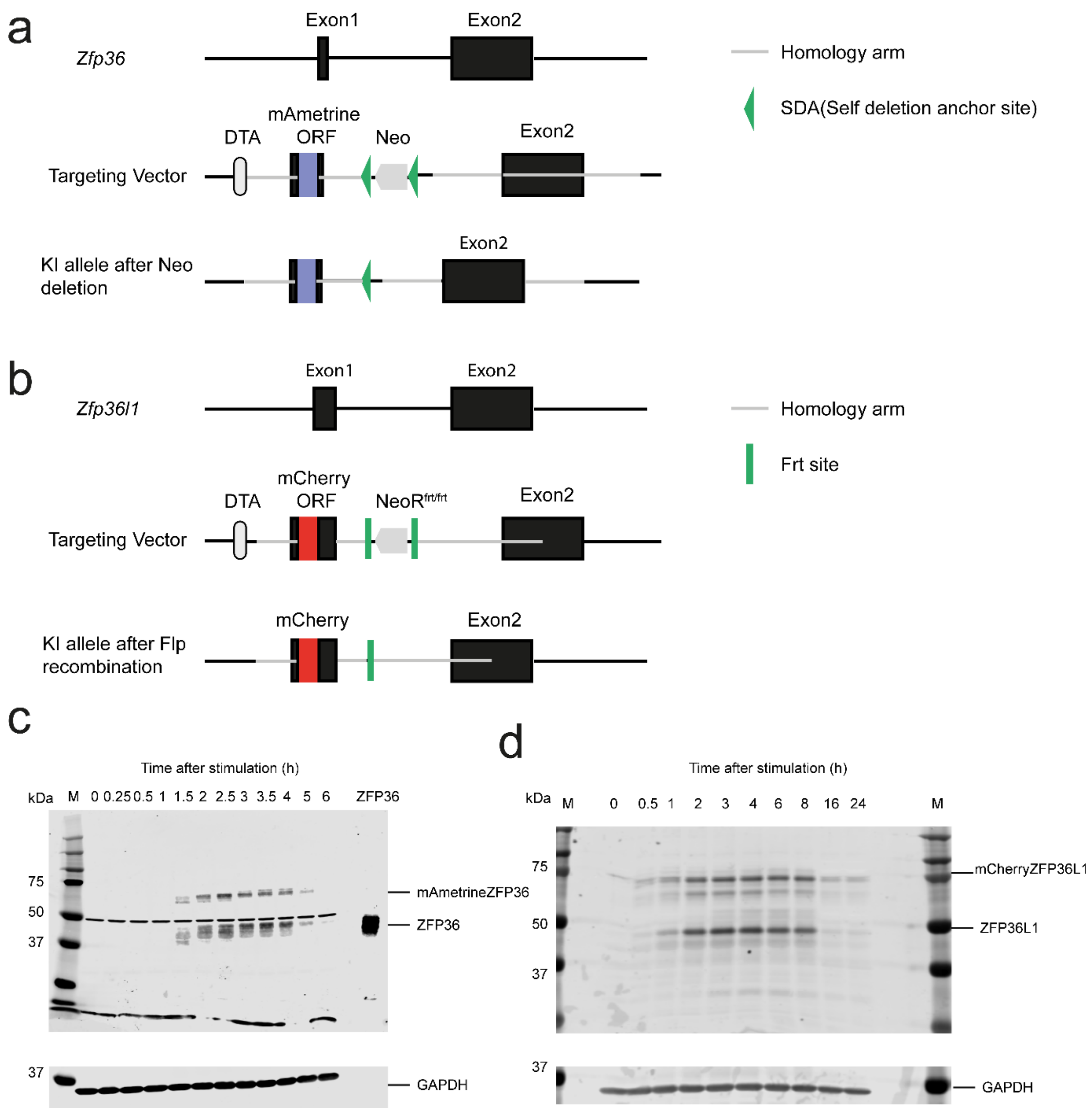


**Supplementary Figure 4: ZFP36 and ZFP36L1 fluorescent reporter mice. a)** Schematic illustrates the targeting vector and the resulting engineered $^{mAmetrine}$ZFP36 allele and $^{mCherry}$ZFP36L1 **b)** allele where the respective gene sequences encoding the fluorescent reporter proteins have been knocked into the first exons of *Zfp36* and *Zfp36l1*. Peptide sequences with highlighted fluorescent reporters are indicated for each fusion protein. Detection of endogenous and fusion protein expression of ZFP36 and ZFP36L1 from heterozygous mice. **c)** Western blot of ZFP36 and $^{mAmetrine}$ZFP36 and **d)** ZFP36L1 and $^{mCherry}$ZFP36L1 expression in *in vitro* expanded CD8 T cells from heterozygous mice stimulated with $10^{-10}$M N4 peptide for the indicated times. Equal amounts of protein were loaded per lane. Last lane in **c)** was loaded with lysate from HEK293 cells expressing ZFP36. Each blot is a representative of at least three independent experiments.
